# RAVEN 2.0: a versatile toolbox for metabolic network reconstruction and a case study on *Streptomyces coelicolor*

**DOI:** 10.1101/321067

**Authors:** Hao Wang, Simonas Marcišauskas, Benjamín J Sánchez, Iván Domenzain, Daniel Hermansson, Rasmus Agren, Jens Nielsen, Eduard J Kerkhoven

## Abstract

RAVEN is a commonly used MATLAB toolbox for genome-scale metabolic model (GEM) reconstruction, curation and constraint-based modelling and simulation. Here we present RAVEN Toolbox 2.0 with major enhancements, including: (i) *de novo* reconstruction of GEMs based on the MetaCyc pathway database; (ii) a redesigned KEGG-based reconstruction pipeline; (iii) convergence of reconstructions from various sources; (iv) improved performance, usability, and compatibility with the COBRA Toolbox. Capabilities of RAVEN 2.0 are here illustrated through *de novo* reconstruction of GEMs for the antibiotic-producing bacterium *Streptomyces coelicolor*. Comparison of the automated *de novo* reconstructions with the iMK1208 model, a previously published high-quality *S. coelicolor* GEM, exemplifies that RAVEN 2.0 can capture most of the manually curated model. The generated *de novo* reconstruction is subsequently used to curate iMK1208 resulting in Sco4, the most comprehensive GEM of *S. coelicolor*, with increased coverage of both primary and secondary metabolism. This increased coverage allows the use of Sco4 to predict novel genome editing targets for optimized secondary metabolites production. As such, we demonstrate that RAVEN 2.0 can be used not only for *de novo* GEM reconstruction, but also for curating existing models based on up-to-date databases. Both RAVEN 2.0 and Sco4 are distributed through GitHub to facilitate usage and further development by the community.

**Author summary:** Cellular metabolism is a large and complex network. Hence, investigations of metabolic networks are aided by *in silico* modelling and simulations. Metabolic networks can be derived from whole-genome sequences, through identifying what enzymes are present and connecting these to formalized chemical reactions. To facilitate the reconstruction of genome-scale models of metabolism (GEMs), we have developed RAVEN 2.0. This versatile toolbox can reconstruct GEMs fast, through either metabolic pathway databases KEGG and MetaCyc, or from homology with an existing GEM. We demonstrate RAVEN’s functionality through generation of a metabolic model of *Streptomyces coelicolor*, an antibiotic-producing bacterium. Comparison of this *de novo* generated GEM with a previously manually curated model demonstrates that RAVEN captures most of the previous model, and we subsequently reconstructed an updated model of *S. coelicolor*: Sco4. Following, we used Sco4 to predict promising targets for genetic engineering, which can be used to increase antibiotic production.

## Introduction

Genome-scale metabolic models (GEMs) are comprehensive *in silico* representations of the complete set of metabolic reactions that take place in a cell [1]. GEMs can be used to understand and predict how organisms react to variations on genetic and environmental parameters [2]. Recent studies demonstrated the extensive applications of GEMs in discovering novel metabolic engineering strategies [3]; studying microbial communities [4]; finding biomarkers for human diseases and personalized and precision medicines [5,6]; and improving antibiotic production [7]. With the increasing ease of obtaining whole-genome sequences, significant challenges remain to translate this knowledge to high-quality GEMs [8].

To meet the increasing demand of metabolic network modelling, the original RAVEN (<u>R</u>econstruction, <u>A</u>nalysis and <u>V</u>isualization of M<u>e</u>tabolic <u>N</u>etworks) toolbox was developed to facilitate GEM reconstruction, curation, and simulation [9]. In addition to facilitating the analysis and visualization of existing GEMs, RAVEN particularly aimed to assist semi-automated draft model reconstruction, utilizing existing template GEMs and the KEGG database [10]. Since publication, RAVEN has been used in GEMs reconstruction for a wide variety of organisms, ranging from bacteria [11], archaea [12] to human gut microbiome [13], eukaryotic microalgae [14], parasites [15–17], and fungi [18], as well as various human tissues [19,20] and generic mammalians models with complex metabolism [21,22]. As such, the RAVEN toolbox has functioned as one of the two major MATLAB-based packages for constraint-based metabolic modelling, together with the COBRA Toolbox [23–25].

Here, we present RAVEN 2.0 with greatly enhanced reconstruction capabilities, together with additional new features (Fig 1, Table 1). A prominent enhancement of RAVEN 2.0 is the use of the MetaCyc database in assisting draft model reconstruction. MetaCyc is a pathway database that collects only experimentally verified pathways with curated reversibility information and mass-balanced reactions [26]. RAVEN 2.0 can leverage this high-quality database to enhance the GEM reconstruction process. While the functionality of the original RAVEN toolbox was illustrated by reconstructing a GEM of *Penicillium chrysogenum* [9], we here demonstrate the new and improved capabilities and wide applicability of RAVEN 2.0 through reconstruction of a GEM for *Streptomyces coelicolor*.

**Table 1.**
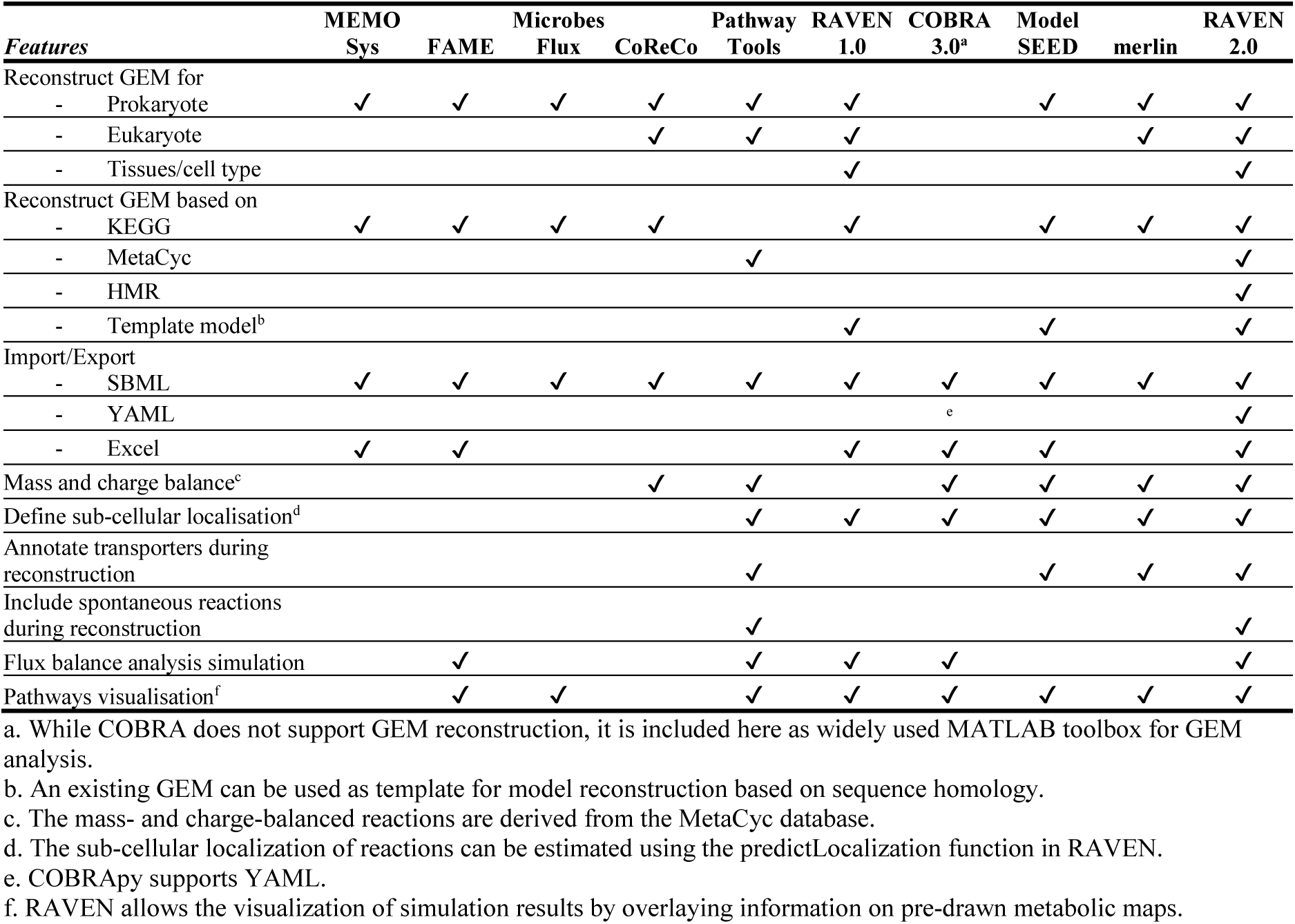
Feature comparison of GEM reconstruction toolboxes.

**Fig 1.**
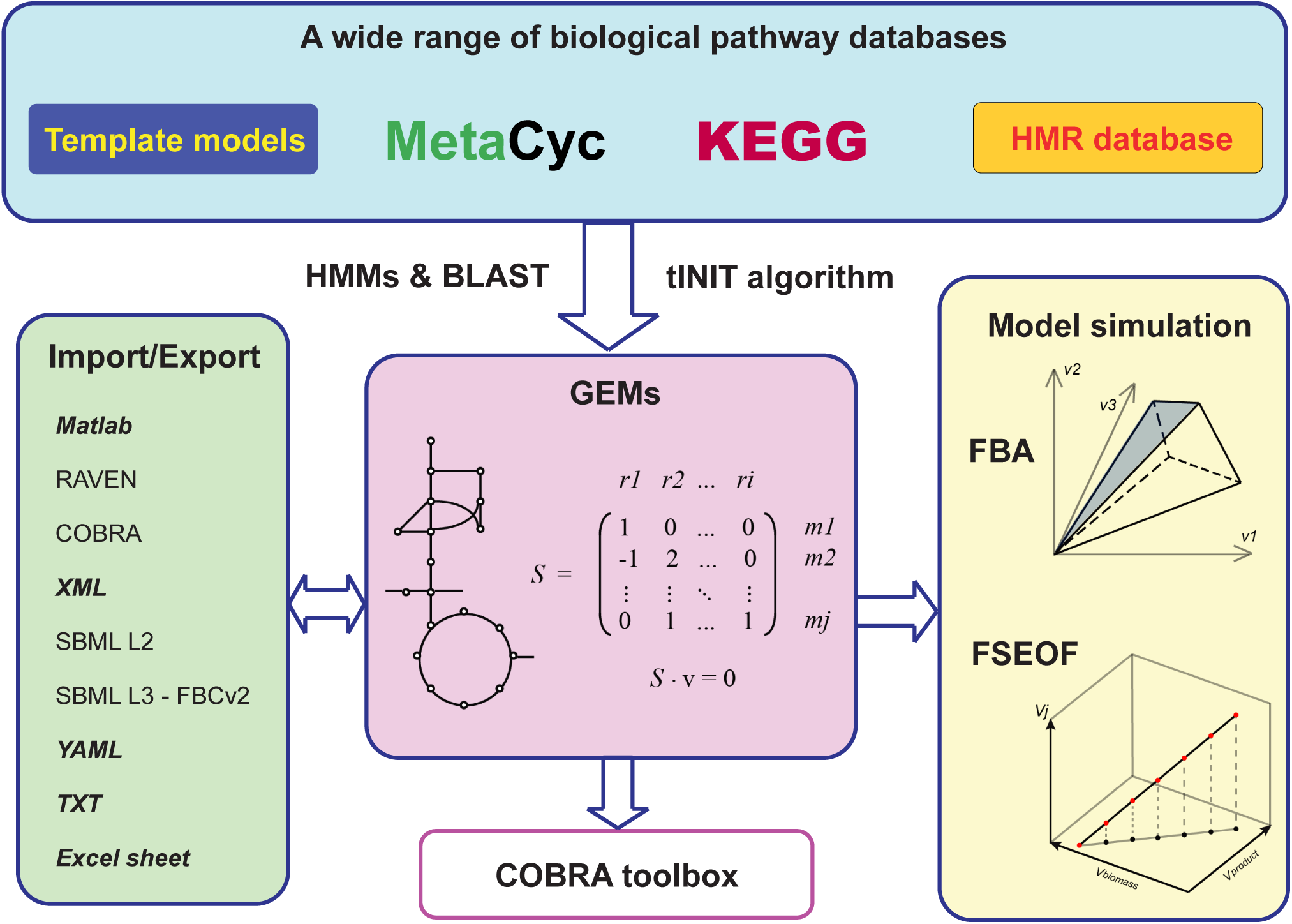
Schematic overview of RAVEN Toolbox version 2.0. RAVEN was significantly enhanced for *de novo* draft model reconstruction by integrating knowledge from different sources (e.g. KEGG and MetaCyc). It has better import/export support to relevant formats, and especially improved compatibility with the COBRA Toolbox by resolving previously conflicting function names and providing a bi-directional model conversion function. As a MATLAB toolbox, RAVEN provides one unified environment for both model reconstruction and simulation analysis (e.g. FSEOF) and allows scripting for more flexible operations.

*S. coelicolor* is a representative species of soil-dwelling, filamentous and Gram-positive actinobacterium harbouring enriched secondary metabolite biosynthesis gene clusters [27,28]. As a well-known pharmaceutical and bioactive compound producer, *S. coelicolor* has been exploited for antibiotic and secondary metabolite production [29]. The first published GEM for *S. coelicolor*, iIB711 [30], was improved through an iterative process resulting in the GEMs iMA789 [31] and iMK1208 [32]. The most recent GEM, iMK1208, is a high-quality model that includes 1208 genes and 1643 reactions and was successfully used to predict metabolic engineering targets for increased production of actinorhodin [32].

Here, we demonstrate how the new functions of RAVEN can be used for *de novo* reconstruction of a *S. coelicolor* GEM, using comparison to the existing high-quality model iMK1208 as benchmark. The use of three distinct *de novo* reconstruction approaches enabled capturing most of the existing model, while complementary reactions found through the *de novo* reconstructions gave the opportunity to improve the existing model. After manual curation, we included 402 new reactions into the GEM, with 320 newly associated enzyme-coding genes, including a variety of biosynthetic pathways for known secondary metabolites (e.g. 2-methylisoborneol, albaflavenone, desferrioxamine, geosmin, hopanoid and flaviolin dimer). The updated *S. coelicolor* GEM is released as Sco4, which can be used as an upgraded platform for future systems biology research on *S. coelicolor* and related species.

## Results

### Genome-scale reconstruction and curation with RAVEN 2.0

RAVEN 2.0 aims to provide a versatile and efficient toolbox for metabolic network reconstruction and curation (Fig 1). In comparison to other solutions for GEM reconstruction (Table 1), the strength of RAVEN is its ability of semi-automated reconstruction based on published models, KEGG and MetaCyc databases, integrating knowledge from diverse sources. A brief overview of RAVEN capabilities is given here, while more technical details are stated in Material & Methods, and detailed documentation is provided for individual functions in the RAVEN package.

RAVEN supports two distinct approaches to initiate GEM reconstruction for an organism of interest: (i) based on protein homology to an existing template model, or (ii) *de novo* using reaction databases. The first approach requires a high-quality GEM of a phylogenetically closely related organism, and the functions *getBlast* and *getModelFromHomology* are used to infer homology using bidirectional BLASTP and build a subsequent draft model. Alternatively, *de novo* reconstruction can be based on two databases: KEGG and MetaCyc. For KEGG-based reconstruction, the user can deploy *getKEGGModelForOrganism* to either rely on KEGG-supplied annotations—KEGG currently includes over 5000 genomes—or query its protein sequences for similarity to HMMs that are trained on genes annotated in KEGG. MetaCyc-based reconstruction can be initiated with *getMetaCycModelForOrganism* that queries protein sequences with BLASTP for homology to enzymes curated in MetaCyc, while *addSpontaneous* retrieves relevant non-enzyme associated reactions from MetaCyc.

Regardless of which (combination of) approach(es) is followed, a draft model is obtained that requires further curation to result in a high-quality reconstruction suitable for simulating flux distributions. Various RAVEN functions aid in this process, including *gapReport* that runs a gap analysis and reports e.g. dead-end reactions and unconnected subnetworks that indicate missing reactions and gaps in the model, in addition to reporting metabolites that can be produced or consumed without in- or output from the model, which is indicative of unbalanced reactions. RAVEN is distributed with a gap-filling algorithm *gapFill*, however, results from external gap-filling approaches can also be readily incorporated. This and further manual curation is facilitated through functions such as *addRxnsGenesMets* that moves reactions from a template to a draft model, *changeGeneAssoc* and *standardizeGrRules* that curate gene associations and *combineMetaCycKEGGModels* that can semi-automatically unify draft models reconstructed from different databases.

In addition to model generation, RAVEN includes basic simulation capabilities including flux balance analysis (FBA), random sampling of the solution space [33] and flux scanning with enforced objective function (FSEOF) [34]. Models can be handled in various file-formats, including the community standard SBML L3V1 FBCv2 that is compatible with many other constraint-based modelling tools, including the COBRA Toolbox [23], as well as non-MATLAB tools as COBRApy [35] and SBML-R [36]. In addition, a user-friendly Excel format is implemented; while support for flat-text and YAML formats are provided to facilitate tracking changes between model versions, which is unfeasible with the SBML file format. As a MATLAB package, RAVEN gives users flexibility to build their own reconstruction and analysis pipelines according to their needs.

### Draft model reconstruction for *S. coelicolor*

The enriched capabilities of RAVEN 2.0 were evaluated by *de novo* generation of GEMs for *S. coelicolor* using three distinct approaches, as described in Material & Methods (Fig 2). Cross-comparison of genes from *de novo* reconstructions and the published *S. coelicolor* GEM iMK1208 indicated that the three *de novo* approaches are complementary and comprehensive, combined covering 88 % of the genes included in iMK1208 (Fig 3). The existing model contained 146 genes that were not annotated by any of the automated approaches, signifying the valuable manual curation that has gone into previous GEMs of *S. coelicolor*. Nonetheless, matching of metabolites across models through their KEGG identifiers further supported that most of the previous GEM is captured by the three *de novo* reconstructions, while each approach has their unique contribution (Fig 3).

**Fig 2.**
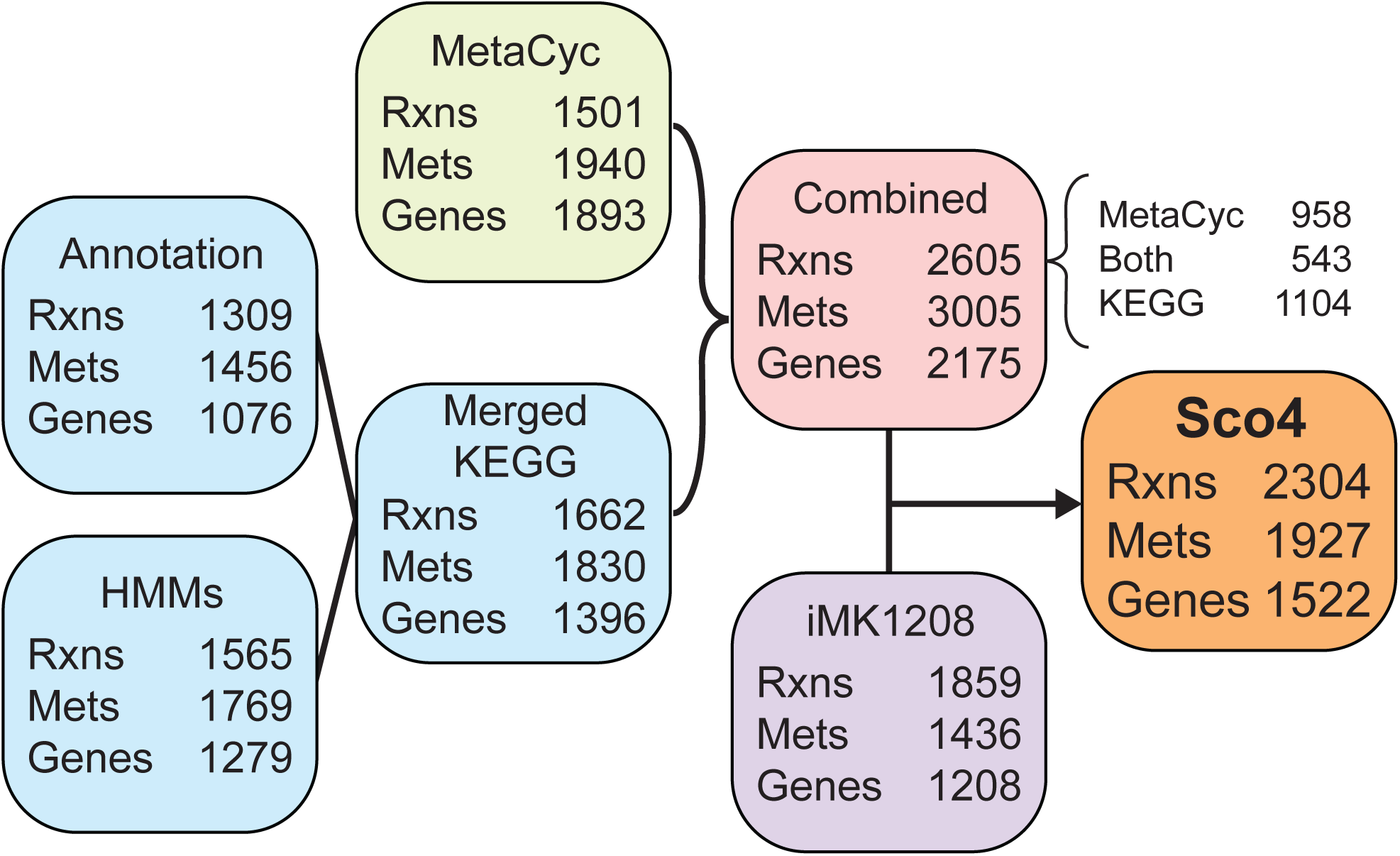
Schematic illustration of Sco4 model generation by RAVEN 2.0. Three draft GEMs were automatically reconstructed in *de novo* based on MetaCyc and KEGG databases. The two GEMs derived from KEGG genome annotation and HMMs were merged and then subsequently combined with the MetaCyc-based model. The reactions and annotations in the combined draft model were used for manual curation of iMK1208 to result in Sco4.

**Fig 3.**
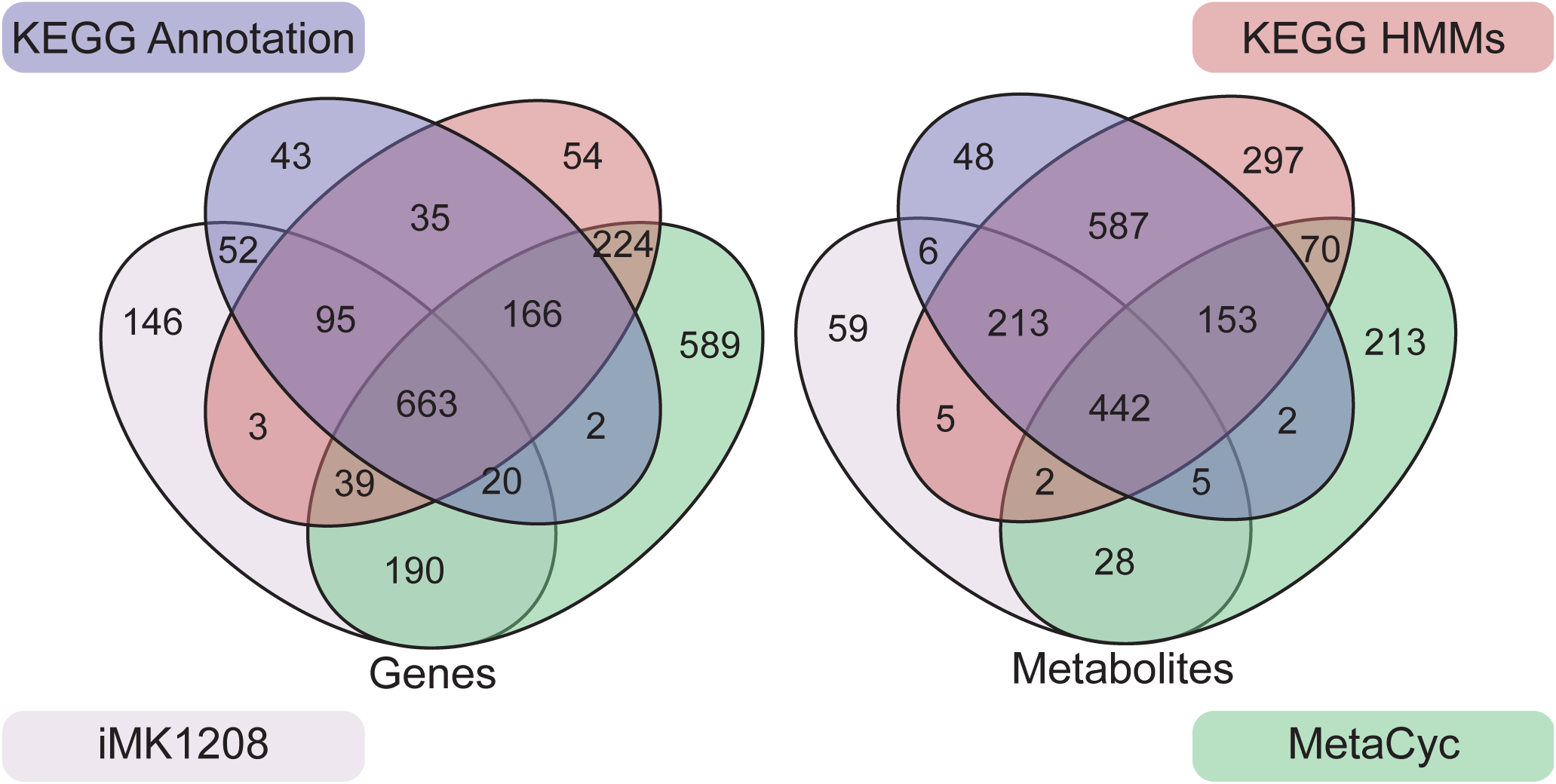
Venn diagrams comparing genes and metabolites between the three *de novo* draft reconstructions and iMK1208.

The three draft reconstructions were consecutively merged to result in a combined draft reconstruction (S1 Data), containing 2605 reactions, of which 958 and 1104 reactions were uniquely from MetaCyc- and KEGG-based reconstructions, respectively (Fig 2). While MetaCyc-based reconstruction annotated more genes than KEGG-based reconstructions (Fig 3), the number of unique reactions by MetaCyc is slightly lower than by KEGG, indicating that KEGG based reconstruction is more likely to assign genes to multiple reactions. Of the 789 reactions from the existing high-quality model that could be mapped to either MetaCyc or KEGG reactions, 733 (92.9%) were included in the combined draft model (S1 Table).

### Further development of the obtained model

The combined *de novo* reconstruction has a larger number of reactions, metabolite and genes than the previous *S. coelicolor* GEM (Fig 2). While a larger metabolic network does not necessarily imply a *better* network, we took advantage of the increased coverage of the *de novo* reconstruction by using it to curate iMK1208, while retaining the valuable contributions from earlier GEMs. The culminating model is called Sco4, the fourth major release of *S. coelicolor* GEM. Through manual curation, a total of 398 metabolic reactions were selected from the combined model to expand the stoichiometric network of the previous GEM (S3 Table). These new reactions cover diverse subsystems including both primary and secondary metabolism (Fig 4A) and displayed close association with existing metabolites in the previous GEM (Fig 4B). Despite both MetaCyc- and KEGG-based reconstructions contributing roughly equally, MetaCyc-unique reactions are more involved in energy and secondary metabolism, while KEGG-unique reactions are more related to amino acid metabolism and degradation pathways (Fig 4C). The *de novo* reconstruction annotated genes to 11 reactions that had no gene association in the previous GEM (S4 Table). Together with 34 spontaneous reactions and 10 transport reactions identified by the MetaCyc reconstruction functions (S5 and S6 Tables), the resulting Sco4 model contains 2304 reactions, 1927 metabolites and 1522 genes (Fig 2).

**Fig 4.**
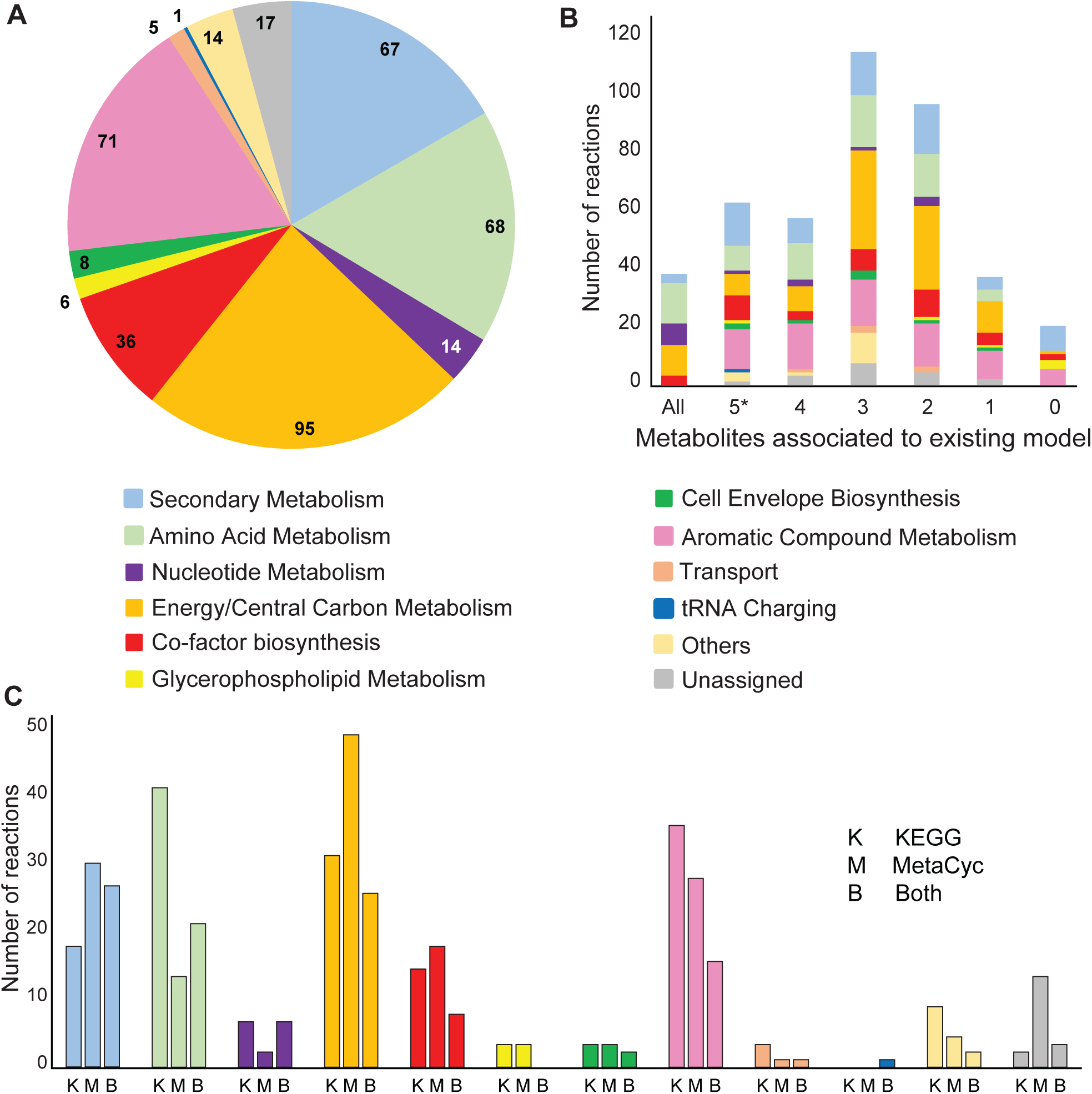
The compositional subsystem distribution of the new reactions selected from the combined draft model. (A) A total of 398 metabolic reactions were introduced in Sco4, thereby expanding the coverage in both primary and secondary metabolism. (B) These 398 curated reactions were identified through either KEGG- and/or MetaCyc-based *de novo* reconstructions. (C) Metabolites from these new reactions are closely associated with the previous *S. coelicolor* GEM. Over 95% of the reactions have one or more metabolites included in iMK1208, and 36 reactions have all reactants and products included in the previous model, resulting in increased connectivity of the network; while only 19 reactions have all newly introduced metabolites, representing completely new stoichiometric network reconstructed by RAVEN 2.0.

The process of model curation using *de novo* reconstructions furthermore identified erroneous annotations in the previous GEM. Seventeen metabolites were annotated with invalid KEGG identifiers (S7 Table), impeding matching with the KEGG-based reconstructions. However, by annotating the reactions and metabolites to MetaCyc, we were still able to annotate all 17 metabolites with a valid KEGG identifier, using MetaCyc-provided KEGG annotations. While the KEGG identifiers used in iMK1208 were valid previously, they have since been removed from the KEGG database. Unfortunately, no changelogs are available to trace such revisions.

### Simulations and predictions with Sco4

The quality of Sco4 was evaluated through various simulations. It displayed the same performance as iMK1208 in growth prediction on 64 different nutrient sources, with a consistent sensitivity of 90.6% (S8 Table). Experimentally measured growth rates in batch and chemostat cultivations were in good correlation with the growth rates predicted by Sco4 (Fig 5A).

**Fig 5.**
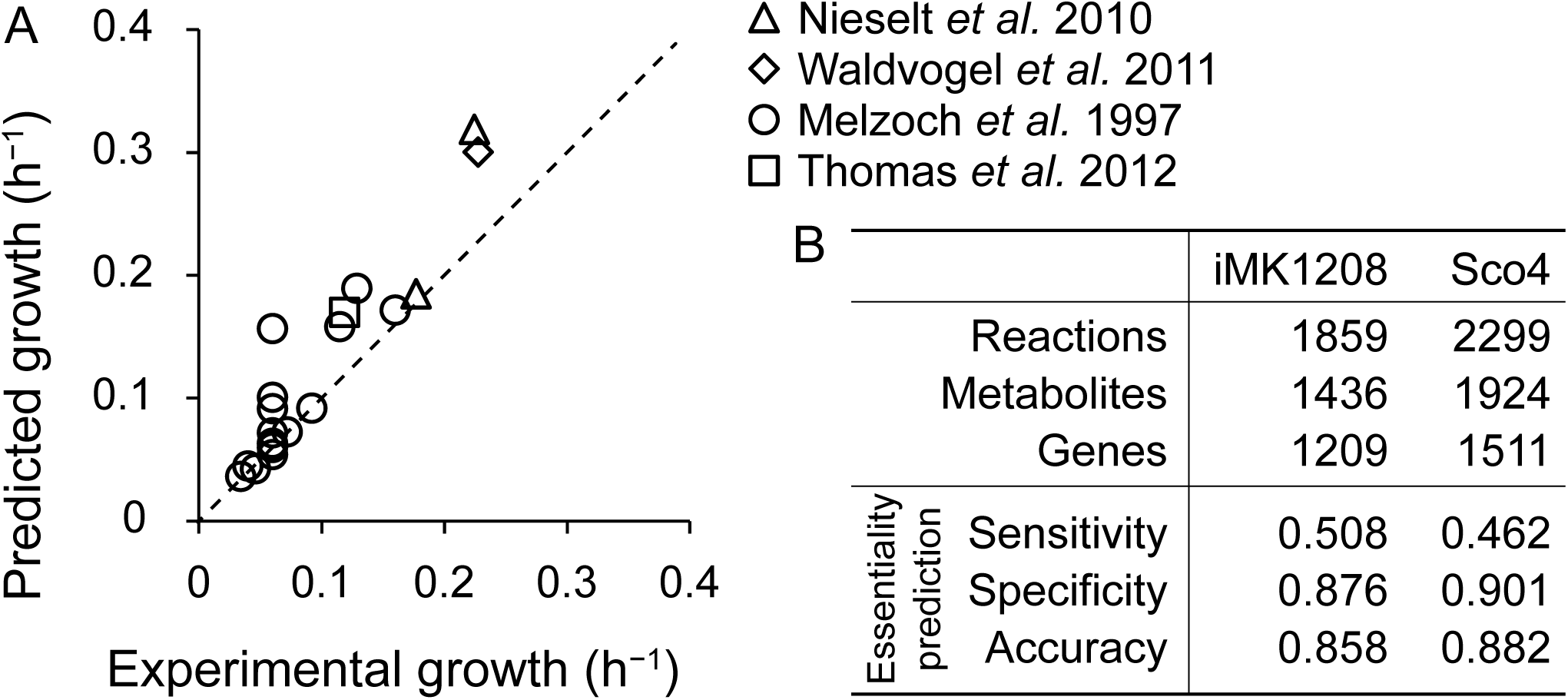
Growth and gene essentiality prediction of Sco4. (A) Comparison of experimental growth rate and predicted growth rate of published bioreactor cultivations [37], using reported exchange rates as model constraints. (B) Comparison between the previously curated GEM iMK1208 and Sco4, of network characteristics and results from single-gene non-essentiality predictions, based on a mutagenesis study by [38].

A recent large-scale mutagenesis study produced and analyzed 51,443 *S. coelicolor* mutants, where each mutant carried a single Tn5 transposition randomly inserted in the genome [37]. No transposition insertions were detected in 79 so-called cold regions of the genome, harboring 132 genes of which 65 are annotated to reactions in Sco4 (S9 Table). The 132 genes are potentially essential, as insertions into these loci would have resulted in a lethal phenotype. However, as it is unclear whether gene essentiality is truly the cause behind the cold-regions, we therefore take the more conservative assumption that genes located *outside* cold regions are *not essential* and compared the non-essential gene sets. Simulation with Sco4 indicates a specificity (or true negative rate) of 0.901, which is an increase over the 0.876 of the previous model (Fig 5B).

The *S. coelicolor* genome project revealed a dense array of secondary metabolite gene clusters both in the core and arms of the linear chromosome (Bentley et al. 2002), and extensive efforts have been made to elucidate these biosynthetic pathways (Van Keulen and Dyson, 2014). The previous GEM of *S. coelicolor* included only three of these pathways (i.e. actinorhodin, calcium-dependent antibiotic and undecylprodigiosin). Through our *de novo* reconstruction, we captured the advances that have since been made in elucidating additional pathways: Sco4 describes the biosynthetic pathways of 6 more secondary metabolites (e.g. geosmin). These additional pathways were mainly obtained from the MetaCyc-based reconstruction (Fig 4C).

The expanded description of secondary metabolism was used to predict potential metabolic engineering targets for efficient antibiotic production in *S. coelicolor*. Flux scanning with enforced objective function (FSEOF) [34] was applied to all secondary metabolic pathways in Sco4 and suggested overexpression targets were compared, with significant overlap between different classes of secondary metabolites (Fig 6, S10 Table). In addition, several targets were predicted to increase production of all modelled secondary metabolites. Three reactions, constituting the pathway from histidine to N-formimidoyl-L-glutamate, and catalyzed by SCO3070, SCO3073 and SCO4932, were commonly identified as potential targets (S10 Table).

**Fig 6.**
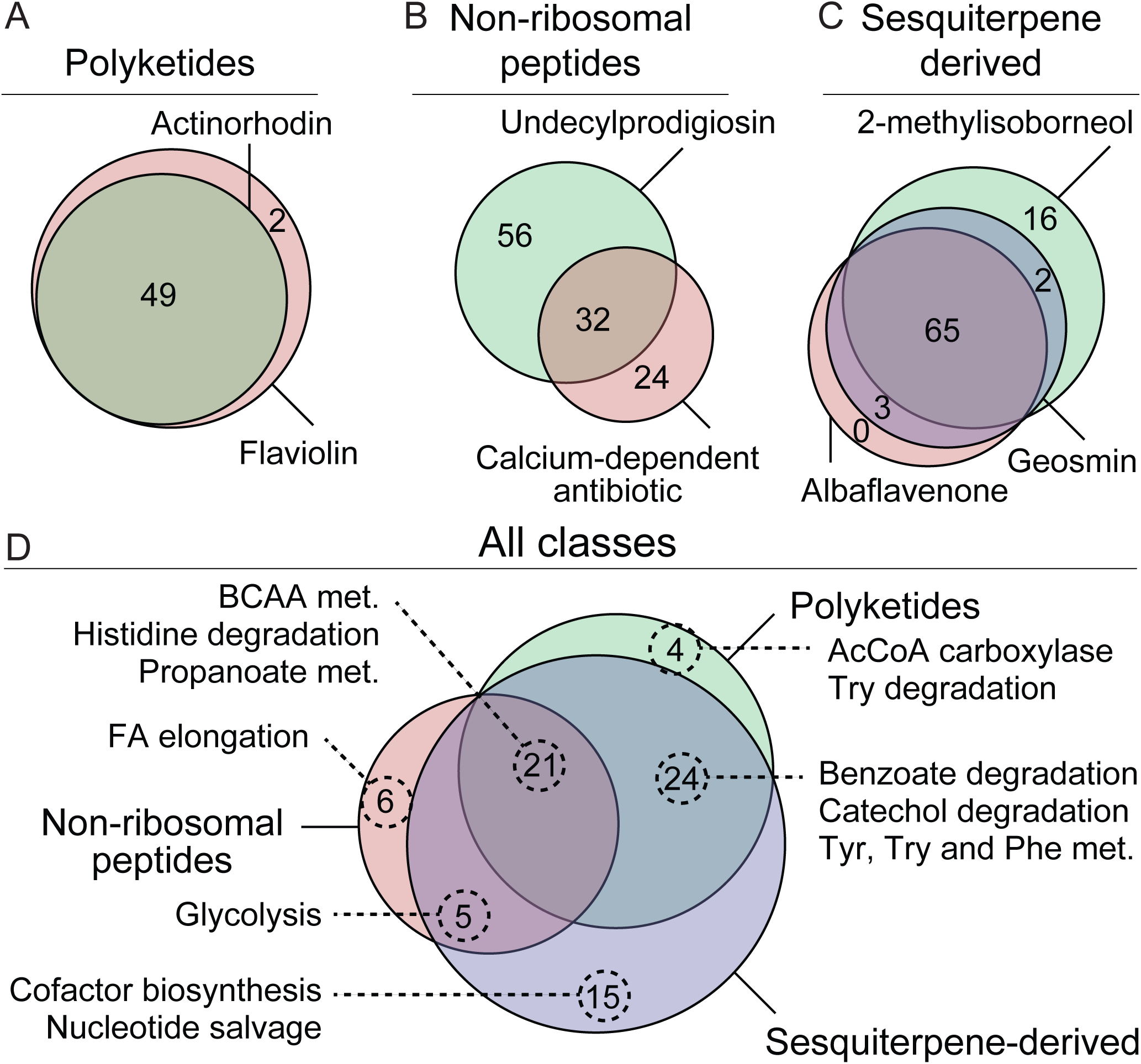
Venn diagrams of targets predicted to be beneficial for overproduction of various secondary metabolites by FSEOF. The 7 secondary metabolites that Sco4 can produce are grouped (A-C), and the overlap of target reactions are indicated within each group. Reactions that overlap within each group are subsequently compared between all groups (D), including in which pathways these reactions are involved. BCAA: branch-chain amino acid; TPI: triosephosphate isomerase; MEP/DOXP: mevalonate independent pathway.

## Discussion

The RAVEN toolbox aims to assist constraint-based modeling with a focus on network reconstruction and curation. A growing number of biological databases have been incorporated for automated GEM reconstruction (Figure 1). The generation of tissue/cell type-specific models through task-driven model reconstruction (tINIT) has been incorporated to RAVEN 2.0 as built-in resource for human metabolic modeling [19,39]. RAVEN 2.0 was further expanded in this study by integrating the MetaCyc database, including experimentally elucidated pathways, chemically-balanced reactions, as well as associated enzyme sequences (21). This key enhancement brings new features toward high-quality reconstruction, such as inclusion of transport and spontaneous reactions (Table 1).

The performance of RAVEN 2.0 in *de novo* reconstruction was demonstrated by the large overlap of reactions between the automatically obtained draft model of *S. coelicolor* and the manually curated iMK1208 model [32]. This indicates that *de novo* reconstruction with RAVEN is an excellent starting point towards developing a high-quality model, while a combined *de novo* reconstruction can be produced within hours on a personal computer. We used the *de novo* reconstructions to curate the existing iMK1208 model, and the resulting Sco4 model was expanded with numerous reactions, metabolites and genes, in part representing recent progress in studies on metabolism of *S. coelicolor* and related species (Figure 5). We have exploited this new information from biological databases to predict novel targets for metabolic engineering toward establishing *S. coelicolor* as a potent host for a wide range of secondary metabolites (Figure 7). Therefore, RAVEN 2.0 can be used not only for *de novo* reconstruction but also model curation and continuous update, which would be necessary for a published GEM to synchronize with the incremental knowledge. We thus deposited the Sco4 as open GitHub repositories for collaborative development with version control.

**Fig 7.**
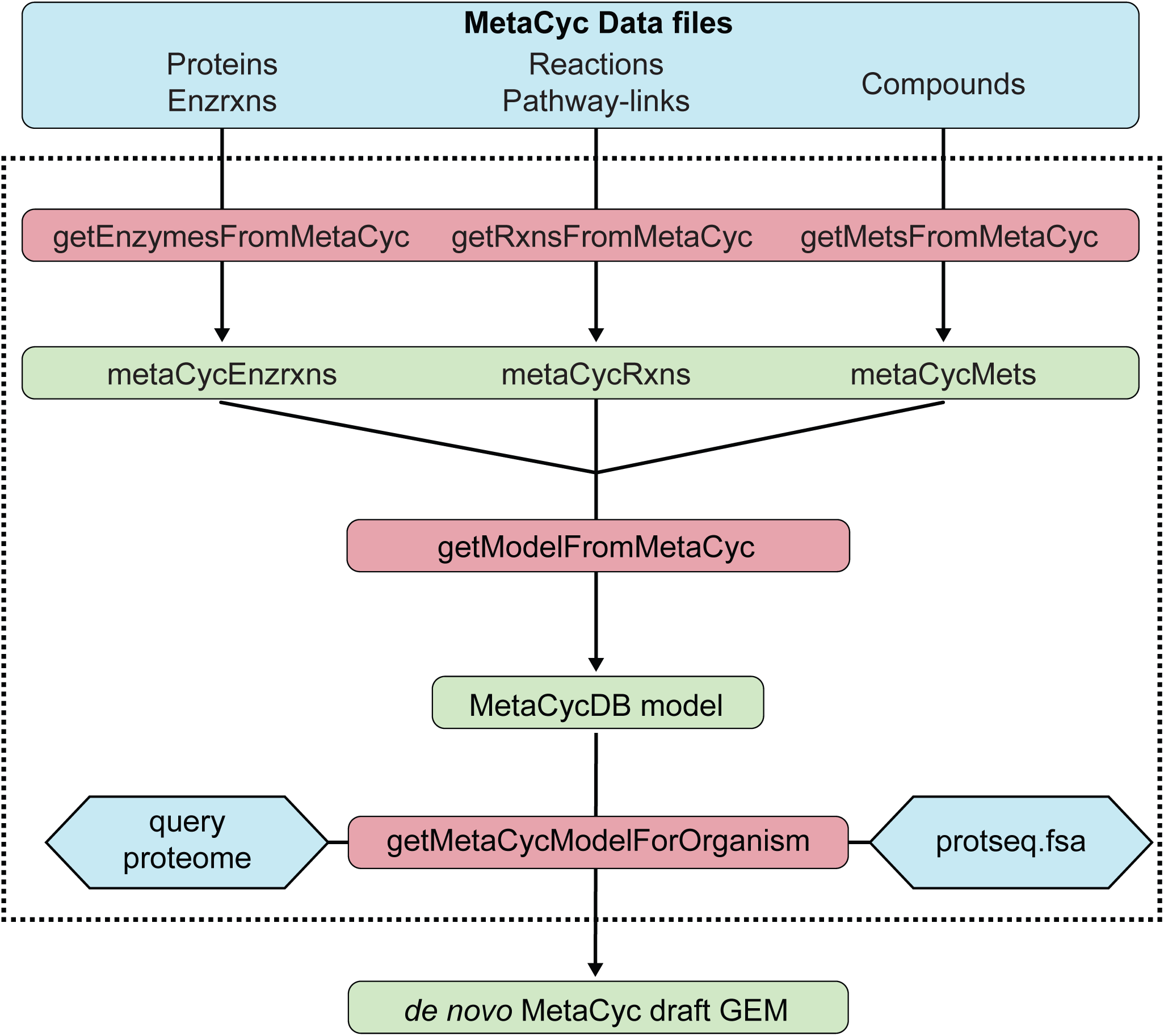
Flow-chart of the functions for MetaCyc-based *de novo* GEM reconstruction in RAVEN 2.0. The MetaCyc-based module (indicated by the dashed line) reads in data files of MetaCyc pathways, reactions, compounds and enzymes, and converts the information into a model structure, which can then be utilized for automatic draft GEM generation through querying the protein sequences in MetaCyc (protseq.fsa). Different components are color coded as MetaCyc data files in blue, MATLAB functions in red, and data structures in green.

While RAVEN 2.0 addresses several obstacles and significantly improves GEM reconstruction and curation, a number of challenges remain to be resolved. One major obstacle encountered is matching of metabolites, whether by name or identifier (e.g. KEGG, MetaCyc, ChEBI). Incompatible metabolite nomenclature, incomplete and incorrect annotations all impede fully automatic matching and rather requires intensive manual curation, especially when comparing and combining GEMs from different sources. Efforts have been made to address these issues, e.g. by simplifying manual curation using modelBorgifier [40]. Particularly worth noting is MetaNetX [41], where the MNXref namespace aims to provide a comprehensive cross reference between metabolite and reactions from a wide range of databases, assisting model comparison and integration. Future developments in this direction ultimately leverage this information to automatically reconcile metabolites and reactions across GEMs. Another major challenge is evaluation and tracking of GEM quality. Here we evaluated Sco4 with growth and gene essentiality simulations (Figure 5, S8 Table, S9 Table), however, the GEM modelling community would benefit from such and additional quality tests according to community standards. Exciting ongoing progress here is memote: an open-source software that is under development that contains a community-maintained, standardized set of metabolic model tests [42]. Given the YAML export functionality in RAVEN already supports convenient tracking of model changes in a GitHub repository, this should ideally be combined with tracking model quality with memote, rendering RAVEN suitable for future GEM reconstruction and curation needs.

## Material and Methods

### RAVEN Toolbox development

The RAVEN Toolbox 1.0 was released as an open-source MATLAB-package [9], that has since seen minor updates and bugfixes. Since 2016, the development of RAVEN has been organized and tracked at a public GitHub repository. This repository provides a platform for the GEM reconstruction community, with users encouraged to report bugs, request new features and contribute to the development.

The RAVEN Toolbox is based on a defined model structure (S11 Table). Design choices dictate minor differences between COBRA and RAVEN structures, however, bi-directional model conversion is supported through *ravenCobraWrapper*. Through resolving previously conflicting function names, RAVEN 2.0 is now fully compatible with the COBRA Toolbox. Detailed documentation on the purpose, inputs and outputs for each function are provided in the *doc* folder.

### MetaCyc-based reconstruction module

Novel algorithms were developed to facilitate *de novo* GEM reconstruction by utilizing the MetaCyc database [26]. In this module, corresponding MATLAB structures were generated from MetaCyc data files (version 21.0) that contained 3118 manually curated pathways with 13,689 metabolites and 15,309 reactions (Fig 7). A total of 17,394 enzymes are associated to these pathways and their protein sequences are included (protseq.fsa). Information from these structures is parsed by *getModelFromMetaCyc* to generate a model structure containing all metabolites, reactions and enzymes. This MetaCyc model can subsequently be used for *de novo* GEM reconstruction through the *getMetaCycModelForOrganism* function (Fig 7). A draft model is generated from MetaCyc enzymes (and associated reactions and metabolites) that show homology to the query protein sequences. Beneficial is that MetaCyc reactions are mass- and charge-balanced, while curated transport enzymes in MetaCyc allow inclusion of transport reactions into the draft model.

In addition, MetaCyc provides 515 reactions that may occur spontaneously. As such reactions have no enzyme association, they are excluded from sequence-based reconstruction and can turn into gaps in the generated models. By cataloguing spontaneous reactions in MetaCyc, the *addSpontaneousRxns* function can retrieve spontaneous reactions depending the presence of the relevant reactants in the draft model.

### KEGG-based reconstruction module

In addition to MetaCyc-based GEM reconstruction, RAVEN 2.0 can utilize the KEGG database for *de novo* GEM reconstruction. The reconstruction algorithms were significantly enhanced in multiple aspects: the reformatted KEGG database in MATLAB format is updated to version 82.0; and the pipeline to train KEGG Orthology (KO)-specific hidden Markov Models is expanded. Orthologous protein sequences, associated to particular KEGG Orthology (KO), are organised into non-redundant clusters with CD-HIT [43]. These clusters are used as input in m-sequence alignment with MAFFT [44], for increased accuracy and speed. The hidden Markov models (HMMs) are then trained for prokaryotic and eukaryotic species with various protein redundancy cut-offs (100%, 90% or 50%) using HMMER3 [45] and can now be automatically downloaded when running *getKEGGModelForOrganism*.

### Combining of MetaCyc- and KEGG-based draft models

To capitalize on the complementary information from MetaCyc- and KEGG-based reconstructions, RAVEN 2.0 facilitates combining draft models from both approaches into one unified draft reconstruction (Fig 2). Prior to combining, reactions shared by MetaCyc- and KEGG-based reconstructions are mapped using MetaCyc-provided cross-references to their respective KEGG counterparts (S12 Table). Additional reactions are associated by *linkMetaCycKEGGRxns* through matching the metabolites, aided by cross-references between MetaCyc and KEGG identifiers (S13 Table). Subsequently, the *combineMetaCycKEGGModels* function thoroughly queries the two models for identical reactions, discarding the KEGG versions while keeping the corresponding MetaCyc reactions. In the combined model, MetaCyc naming convention is preferentially used such that unique metabolites and reactions from KEGG-based draft model are replaced with their MetaCyc equivalents whenever possible. The combined draft model works as a starting point for additional manual curation, to result in a high-quality reconstruction.

### Miscellaneous improvements

RAVEN 2.0 contains a range of additional enhancements. Linear problems can be solved through either the Gurobi (Gurobi Optimization Inc., Houston, Texas) or MOSEK (MOSEK ApS, Copenhagen, Denmark) solvers. Various file formats are supported for import and export of models, including Microsoft Excel through Apache POI (The Apache Software Foundation, Wakefield, Massachusetts), the community standard SBML Level 3 Version 1 FBC Package Version 2 through libSBML [46] and YAML for easy tracking of differences between model files. Meanwhile, backwards compatibility ensures that Excel and SBML files generated by earlier RAVEN versions can still be imported.

### From *de novo* draft GEMs to Sco4

An improved GEM of *S. coelicolor*, called Sco4 for the fourth major published model, was generated through RAVEN 2.0 following the pipeline illustrated in Fig 3. The model is based on the complete genome sequences of *S. coelicolor* A3(2), including chromosome and two plasmids (GenBank accession: GCA_000203835.1) [27]. MetaCyc-based draft model was generated with *getMetaCycModelForOrganism* using default cut-offs (bit-score ≥ 100, positives ≥ 45%). Two KEGG-based draft models were generated with *getKEGGModelForOrganism* by (i) using ‘sco’ as KEGG organism identifier, and (ii) querying the *S. coelicolor* proteome against HMMs trained on prokaryotic sequences with 90% sequence identity. These two models were merged with *mergeModels*, subsequently combined with the MetaCyc-based draft using *combineMetaCycKEGGModels*, followed by manual curation. Reactions were mapped from iMK1208 to MetaCyc and KEGG identifiers in a semi-automated manner (S1 Table). Metabolites in iMK1208 were associated to MetaCyc and KEGG identifiers through examining the mapped reactions (S2 Table). Pathway gaps and invalid metabolite identifiers were thus detected and revised accordingly.

Manual curation of the combined draft and iMK1208 culminated in the Sco4 model. Curation entailed identifying reactions from the combined draft, considering the absence of gene-associations in iMK1208; explicit subsystem and/or pathway information; support from both MetaCyc and KEGG reconstructions; additional literature information, as well as potential taxonomic conflicts. Manual curation was particularly required for secondary metabolite biosynthetic pathways, due to high levels of sequence similarity among the synthetic domains of polyketide synthase and nonribosomal peptide synthetase [47]. The identified new reactions were added to Sco4, while retaining the previous manual curation underlying iMK1208. Spontaneous reactions were added through *addSpontaneousRxns*, while transport reactions annotated in the MetaCyc-based reconstruction were manually curated. Gene essentiality was simulated on iMK1208 and Sco4 by the COBRA function *singleGeneDeletion*, with a more than 75% reduction in growth rate identifying essential reactions. Potential targets for metabolic engineering were predicted using the flux scanning with enforced objective function *FSEOF* [34]. The reconstruction and curation of Sco4 is provided as a MATLAB script in the ComplementaryScripts folder of the Sco4 GitHub repository.

### Model repository

The updated Sco4 model is deposited to a GitHub repository in MATLAB .mat, SBML L3V1 FBCv2 .xml, Excel .xlsx, YAML .yml and flat-text .txt formats. Users can not only download the most recent version of the model, but also report issues and suggest changes. Updates in the metabolic network or gene associations can readily be tracked by querying the difference in the flat-text model and YAML representations. As such, Sco4 aims to be a community model, where improved knowledge and annotation will incrementally and constantly refine the model of *S. coelicolor.*

## Acknowledgements

We thank Dr. Sylvain Prigent for valuable discussions.

## Availability

RAVEN is an open source software package available in the GitHub repository (https://github.com/SysBioChalmers/RAVEN). The updated *S. coelicolor* genome-scale metabolic model Sco4 is available as a public GitHub repository at (https://github.com/SysBioChalmers/Streptomyces_coelicolor-GEM). Supporting information is available from FigShare (https://doi.org/10.6084/m9.figshare.6236903).

## Supporting information

Available from FigShare (https://doi.org/10.6084/m9.figshare.6236903).

**S1 Data. Model structure after merging of the three *de novo* draft reconstructions.** The MetaCyc- and KEGG-based *de novo* reconstructions for *S. coelicolor* are merged using the *combineMetaCycKEGGModels* function, as detailed in the Sco4Reconstruction script available from the Sco4 GitHub repository under ComplementaryScripts.

**S1 Table. Mapping of reactions from iMK1208 to MetaCyc and KEGG identifiers.** To facilitate comparison of the existing iMK1208 model and the *de novo* reconstructions, iMK1208 reactions were mapped to their respective KEGG and MetaCyc identifiers. This table is used in the Sco4Reconstruction script available from the Sco4 GitHub repository under ComplementaryScripts.

**S2 Table. Mapping of metabolites from iMK1208 to KEGG and MetaCyc identifiers.** To facilitate comparison of the existing iMK1208 model and the *de novo* reconstructions, iMK1208 metabolites were mapped to their respective KEGG and MetaCyc identifiers. This table is used in the Sco4Reconstruction script available from the Sco4 GitHub repository under ComplementaryScripts.

**S3 Table. Reactions selected from the *de novo* draft model for generating Sco4.** The merged *de novo* model (S1 Data) was manually evaluated to identified 398 reactions that constitute new pathways absent in iMK1208 and were subsequently added towards Sco4. This table is used in the Sco4Reconstruction script available from the Sco4 GitHub repository under ComplementaryScripts.

**S4 Table. Missing gene-associations in iMK1208 that were identified through *de novo* reconstruction.** Comparison of iMK1208 with the merged *de novo* reconstructions identified reactions that had no gene associated in iMK1208, while RAVEN 2.0 was able to identify the responsible genes. This table is used in the Sco4Reconstruction script available from the Sco4 GitHub repository under ComplementaryScripts.

**S5 Table. Spontaneous reactions included in Sco4, as identified from MetaCyc.** The *retrieveSpontaneous* function was used to identify spontaneous reactions that connect to the Sco4 model, as detailed in the Sco4Reconstruction script available from the Sco4 GitHub repository under ComplementaryScripts.

**S6 Table. Transport reactions added to Sco4 based on MetaCyc-based reconstruction and manual curation.** The merged *de novo* model (S1 Data) was manually evaluated to identified 10 transport reactions that were subsequently added to Sco4. This table is used in the Sco4Reconstruction script available from the Sco4 GitHub repository under ComplementaryScripts.

**S7 Table: Complete list of invalid KEGG identifiers and their replacements.** From iMK1208, 17 invalid KEGG metabolite identifiers were identified. Matching 13 of these metabolites to MetaCyc gave their valid KEGG identifier, leveraging on the annotation that is provided by MetaCyc. For the remaining 4 metabolites, it required matching their reaction between MetaCyc and KEGG to identify valid KEGG metabolite identifiers.

**S8 Table. Comparison of growth media validation results between iMK1208 and Sco4.** Culture and simulation conditions were as described previously (Kim *et al.*, 2014). Indicated are carbon and nitrogen sources that have been experimentally determined to be able to support growth of *S. coelicolor*, their exchange reaction, and whether growth is observed in Sco4 and iMK1208. Predictions on iMK1208 and Sco4 show identical results, indicating that the model expansion did not negatively affect model predictions.

**S9 Table: Identification of putative essential genes from mutagenesis study and predictions from the iMK1208 and Sco4 models.** Indicated is whether a gene has been identified as putative essential based on the Tn5 mutagenesis study (Xu *et al*., 2017) and simulation of gene essentiality in iMK1208 or Sco4.

**S10 Table. Combined FSEOF results on the secondary metabolites included in Sco4 model.** Target reactions identified from Sco4 for each of the indicated secondary metabolism product, as obtained from FSEOF analysis. Direction indicates whether a reaction is reversible and carries a flux during the FSEOF analysis in the forward direction (1), reverse direction (−1), or is irreversible (0).

**S11 Table. Definition of RAVEN model structure fields.** Where applicable, corresponding COBRA model structure fields are indicated.

**S12 Table. Matching of reaction identifiers between KEGG and MetaCyc.** Mapping is based on information provided by MetaCyc, while additional matches were identified by comparing reactants using the *linkMetaCycKEGGRxns* function.

**S13 Table. Matching of metabolite identifiers between KEGG and MetaCyc.** Mapping is based on information provided by MetaCyc.

